# Dissecting current rectification through asymmetric nanopores

**DOI:** 10.1101/2024.11.14.623680

**Authors:** Yichun Lin, Jerome J. Lacroix, James D. Sterling, Yun Lyna Luo

**Affiliations:** Department of Biotechnology and Pharmaceutical Sciences, Western University of Health Sciences, Pomona, CA 91766; Department of Basic Medical Sciences, Western University of Health Sciences, Pomona, CA 91766; Henry E. Riggs School of Applied Life Sciences, Keck Graduate Institute, Claremont, CA, 91711

## Abstract

Rectification, the tendency of bidirectional ionic conductors to favor ion flow in a specific direction, is an intrinsic property of many ion channels and synthetic nanopores. Despite its frequent occurrence in ion channels and its phenomenological explanation using Eyring’s rate theory, a quantitative relationship between the rectified current and the underlying ion-specific and voltage-dependent free energy profile has been lacking. In this study, we designed nanopores in which potassium and chloride current rectification can be manipulated by altering the electrostatic pore polarity. Using molecular dynamics-based free energy simulations, we quantified voltage-dependent changes of free energy barriers in six ion-nanopore systems. Our results illustrate how the energy barriers for inward and outward fluxes become unequal in the presence of an electromotive driving force, leading to varying degrees of rectification for cation and anion currents. This work establishes a direct link between equilibrium potential of mean force and current rectification rate and demonstrates that rectification caused by energy barrier asymmetry depends on the nature of the permeating ion, can be tuned by pore polarity, does not require ion binding sites, conformational flexibility, or specific pore geometry, and, as such, may be widespread among ion channels.

**Statement of significance:** In many ion channels, ions flow faster in one direction than the other, even under equal magnitude of driving forces. This phenomenon, known as rectification, often arises as an intrinsic biophysical property of the channel pore and has significant physiological implications. Our molecular dynamics simulations provide a unified framework to dissect the free energy profiles that underpin current rectification. This approach has broad applications in the field of ion channel biology and material design.

## Introduction

Ion channels are essential for the transport of ions, small molecules, or nutrients within biological systems. When activated by environmental stimuli, these channels regulate the flow of ions, generating inward and outward currents that trigger a cascade of downstream signaling events. The term “current rectification” describes the phenomenon of asymmetric conductance, where the application of equal but opposite voltages across an ion channel results in different magnitude of inward vs. outward currents. Beyond biological channels, understanding the physical mechanisms behind current rectification is important for the design of efficient ionic semiconductor diodes and artificial ion pumps (1-4).

Experimentally observed ion channel rectification arises from diverse mechanisms, including voltage-dependent pore blockage (5), conformational rearrangement (6), and intrinsic biophysical properties of the pore itself (7-11). The last one has been extensively studied in minimalist nanopores, such as the bacterial toxin alpha-hemolysin (12-15), as well as in synthetic nanopores made from organic and inorganic polymers (16-20). These studies generally support the notion that intrinsic rectification is linked to asymmetric charge density along the ion conduction pathway. In synthetic nanopores, this charge asymmetry is often exacerbated using conically-shaped pores (4, 14, 21).Yet, the underlying free energy profile governing the observed rectification has not been delineated.

From a thermodynamic perspective, a unifying mechanism of all current rectification is the voltage-dependent alteration in the free energy barriers of ion permeation (22). However, a direct experimental determination of the shape of these energy barriers is challenging. Thus, these barrier models have primarily been used to phenomenologically describe rectification based on experimental conductance data (23, 24). Computationally, mean-field continuum models, such as various Poisson-Nernst-Planck (PNP) models, have been widely and successfully applied to simulate rectified currents within nanopores (4, 18, 19, 24, 25). Beyond continuum approaches, atomistic molecular dynamics (MD) simulations under voltage have also reported rectified I-V curves in nanopores (13-15, 20). However, in these MD simulations, a direct link between the rectified current and the underlying ion-specific and voltage-dependent free energy profile has not been quantitatively established.

In this study, we employed atomistic MD simulation to compute ion-specific free energy barriers explicitly from the physical properties of the ions and channels under equal but opposite voltage. To this aim, we designed a set of nanopores whose potassium and chloride current rectification can be added or removed at will by manipulating electrostatic pore polarity. Briefly, for each ion-nanopore system, we first computed the steady-state flux to quantify the rectification. We then calculated free energy profile of ion permeation (potential of mean force, or PMF) and the mean first-passage-time (MFPT) under equilibrium conditions (zero voltage). From these equilibrium PMFs, we derived non-equilibrium PMFs under negative and positive voltages.

Our results illustrate quantitatively how the energy barriers of inward and outward flux become unequal in the presence of an electromotive driving force, causing both potassium and chloride currents to rectify, albeit to varying degrees. This establishes a direct correlation between equilibrium PMF and current rectification. Our atomistic MD-generated data does not rely on mean-filed approximations, and hence can be readily applied to complex biological channels, in presence or absence of ion/ligand binding. The detailed information from the equilibrium and non-equilibrium free energy profile will enable us to better understand and design biological and synthetic channels with desired rectification properties.

## Methods

### Nanopore setup and simulation protocols

The coordinates of the neutral nanopore were taken from ref (26). The two charged nanopores were constructed by adding -0.5 or +0.5 charge to the two carbon atoms at the intracellular side of the channel (**Fig. 1a**). These channels have sizes of 13.5 Å in length and 6 Å in radius. Two carbon sheets form an artificial membrane to separate the bulk solution. Constraints were applied to all atoms to keep the system rigid. CHARMM36 force field (27, 28) is used throughout this study. After solvation, the box size was 38×38×75 Å^3^, which contained 2503 TIP3P water molecules. 6 to 7 K^+^ and Cl^-^ ions were added to main the charge neutrality of each system. For all simulations, ions in bulk were confined within a 6 Å radius cylinder with a flat-bottom harmonic restraint (force constant 10 kcal/mol·Å^2^) on the xy-plane. Based on the volume of the cylinder (8478 Å^3^), 6 and 7 ions correspond to the bulk concentration of 1.17 and 1.37 M. Since the PMF value is offset to zero at bulk region, this concentration difference has negligible effect on the PMF barrier (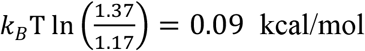 at room temperature 298.15 K).

**Figure 1.**
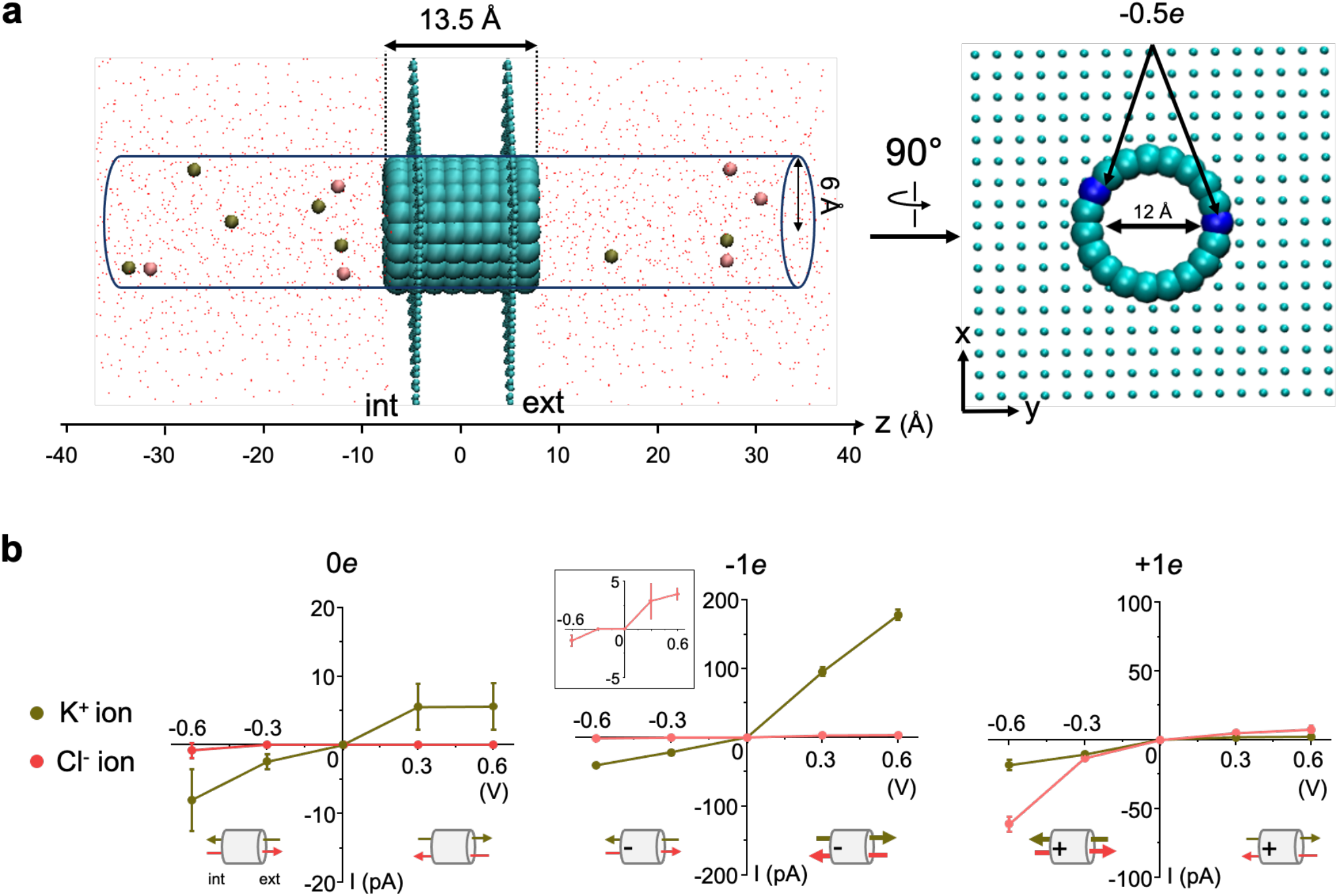
A single charge in cylindrical nanopores rectify ionic currents. (**a**) *Left*: side view of the negatively-charged nanopore system. Nanopore carbon atoms are shown as cyan spheres (vdW representation). The membrane is made from two parallel carbon sheets shown as cyan lines. The 6 Cl^-^ ions and 7 K^+^ ions in the bulk, shown as pink and brown spheres, are confined within a 6 Å radius cylinder. Red dots are oxygen from water molecules. *Right*: nanopore viewed from the intracellular membrane plane showing the two carbon atoms carrying a -0.5*e* charge (blue vdW spheres). (**b**) *I-V* plots for the neutral (0*e*), positively-charged (+1*e*) and negatively-charged (−1*e*) nanopores. By convention, a negative voltage means the intracellular potential is more negative than the extracellular potential. The *I-V* curve for Cl^-^ ions in the -1*e* system is magnified in the insert for clarity. Error bars = standard deviations from two replicas.

Periodic boundary conditions were used, which guarantee that intracellular and extracellular bulk solutions are identical. All MD simulations were performed using NAMD2.13 at constant volume and constant temperature (300 K) using Langevin thermostat. The standard cutoff for calculating vdW interaction and short-range electrostatic interaction was set at 12 Å with force-switched at 10 Å. Long-range electrostatic interactions were calculated using the particle mesh Ewald algorithm (29). The systems underwent a 100 ns equilibrium period before conducting milestoning and voltage simulations.

### Voltage-driven flux and electrostatic potential calculations

After a 100 ns equilibrium simulation, all systems were subjected to ±300 mV and ±600 mV membrane potential perpendicular to the membrane using NAMD2.13. Each voltage setting ran two 200 ns replicas with different initial velocities. By convention, positive currents correspond to the outward (i.e., from the intracellular to the extracellular side) displacement of positively-charged ions. For each voltage, the ion-specific current was computed from the total number of permeation events over the simulation period. The electrostatic potential along the central pore axis in presence and absence of the external voltage (i.e., *ϕ*_*v*_(*z*) and *ϕ*_0_(*z*)) were computed using the VMD PMEPot plugin (30) without the ions inside the pore.

### Markovian Milestoning simulations

The equilibrium potential of mean force (PMF) profile and mean first-passage time (MFPT) for six ion-nanopore systems were computed from Markovian Milestoning simulations following the protocol described in detail in (26, 31). Briefly, the z-coordinate of the tagged ion was used to define a set of Voronoi cells 2 Å apart along the channel pore, and the milestones are the boundaries between the cells. For each nanopore, an average of 36 simulations with the tagged K^+^ or Cl^-^ ion confined in each cell (using a flat-bottom harmonic restraint of force constant 100 kcal/mol·Å^2^) were conducted. During these simulations, the position of the tagged ion was recorded at the frequency of 0.1 ps. The PMF and MFPT profiles were computed from the transition probability matrix and rate matrix according to previous protocol (26, 31). The total simulation time is between 1.0 to 1.5 *μs* for each ion-nanopore system. The convergence is determined when the MFPT for inward and outward permeations reach the same value for each system, as previously discussed (26).

## Results

We start with a symmetric cylindrical nanopore that has been shown to reproduce physiologically relevant channel conductance (26). Unlike biological channels, this nanopore is rigid, contains no ion binding site, and is too short to allow occupancy of multiple ions, eliminating the influence of conformational change, binding site, and ion-ion interactions on our rectification results. To create an asymmetric permeation energy barrier, we introduced -0.5 or +0.5 electronic charge (*e*) to two carbon atoms at one end of the nanopore, arbitrarily defined as the intracellular side, giving the nanopore either a 0*e* (neutral), -1*e* (negative), or +1*e* (positive) charge (a snapshot of the -1*e* nanopore simulation system is illustrated in **Fig. 1a**). We implemented periodic boundary conditions to maintain ionic concentrations symmetric and constant during non-equilibrium steady-state ion flux simulations under ±300 and ±600 mV. From these simulations, we computed current vs. voltage (*I-V*) curves, ionic conductance and rectification scores (*r*) (**Fig. 1b and Table 1**), the latter corresponding to the ratio of positive vs. negative current at the same voltage magnitude (13).

**Table 1.**
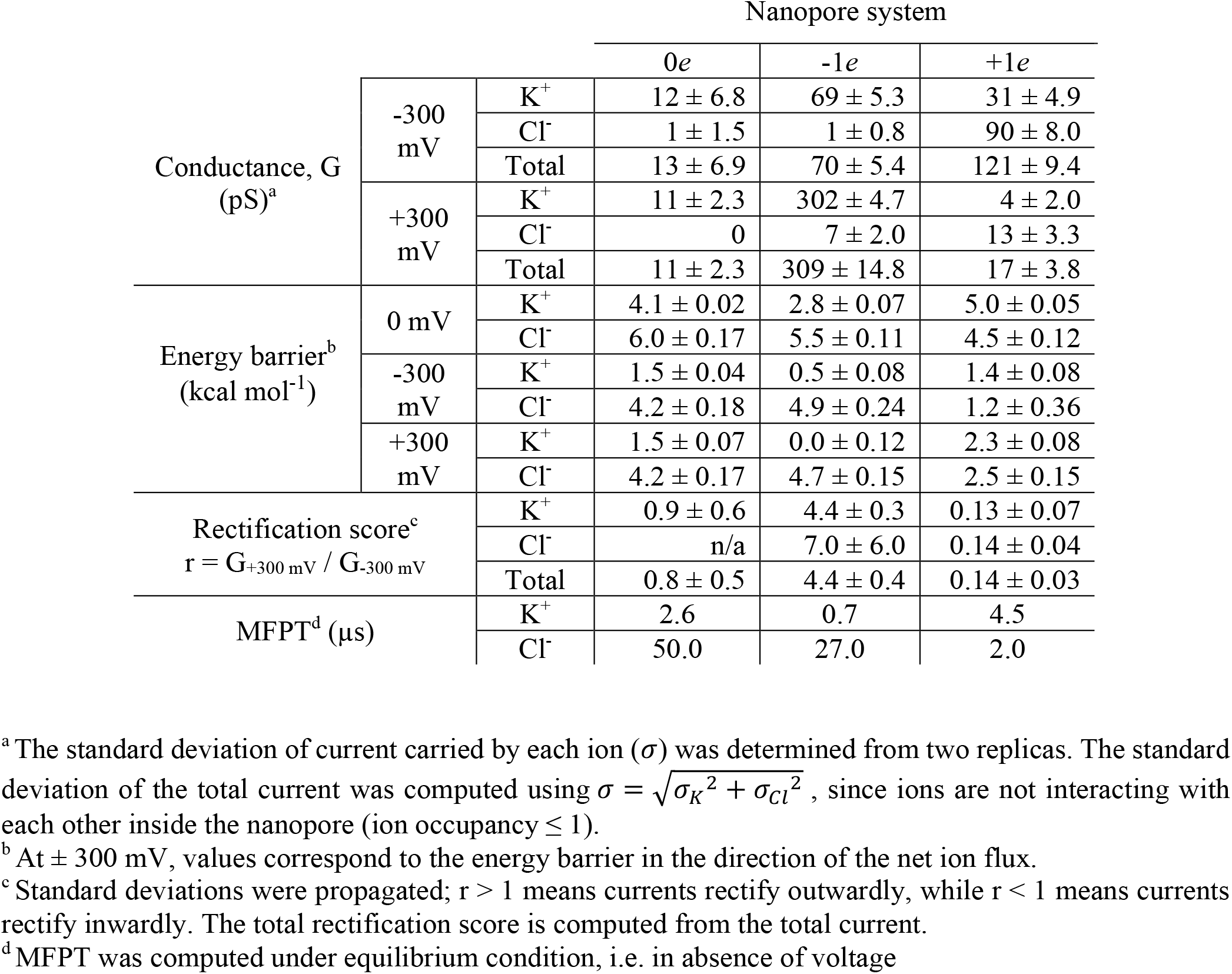
Summary of computational data for each ion-nanopore system.

Our voltage-driven flux simulations show that neither potassium nor chloride currents rectify in the neutral nanopore, as expected since this system is perfectly symmetric. The negative nanopore largely increases potassium conductance, but more so at a positive vs. negative voltage (+V vs. - V), increasing from 11 ± 2.3 to 302 ± 4.7 pS at +V and from 12 ± 6.8 to 69 ± 5.3 pS at -V. Chloride conductance undergoes minimal change at -V, but slightly increases at +V. Hence, both potassium and chloride currents rectify outwardly in the negative nanopore (rectification score r = 4.4 and 7.0, respectively). In contrast, the positive nanopore is weakly (∼3 fold) chloride selective and the conductance of both ions is higher at -V than +V, rectifying currents inwardly (r = 0.13 and 0.14 for K^+^ and Cl^-^ ions, respectively).

Our MD simulations above revealed that ionic currents rectify outwardly in the negative nanopore but inwardly in the positive nanopore. To understand the underlying mechanism of these computationally observed rectification phenomenon, we first computed the equilibrium PMF and MFPT profiles in absence of external driving force for each ion-nanopore system (**Fig. 2a, 2b** and **Table 1**). Under equilibrium conditions, ion displacement through the nanopore occurs through random walk, producing no net flux. Hence, the equilibrium PMF and MFPT determine the intrinsic ionic selectivity and permeability for each system. The selectivity is clearly seen in the equilibrium PMF for the neutral nanopore, which displays a larger energy barrier for Cl^-^ vs. K^+^ ions (**Fig. 2a** blue). The radius of the nanopore is designed to cause partial dehydration of both ions. In CHARMM36 force field with TIP3P water model, the desolvation free energies of K^+^ and Cl^-^ are similar (−79.5 vs. -79.4 kcal/mol), but Cl^-^ has a larger hydration radius than K^+^ (3.85 Å vs. 3.45 Å, **Fig. S1a**). On average, K^+^ loses about one water and Cl^-^ loses about two water molecules when passing the pore, hence the origin of the K^+^ selectivity in neutral pore (**Fig. S1b**). Consequently, in this nanopore, the MFPT is 19-fold longer for a Cl^-^ than for a K^+^ ion (50.0 vs 2.6 µs) (**Fig. 2b** blue).

**Figure 2.**
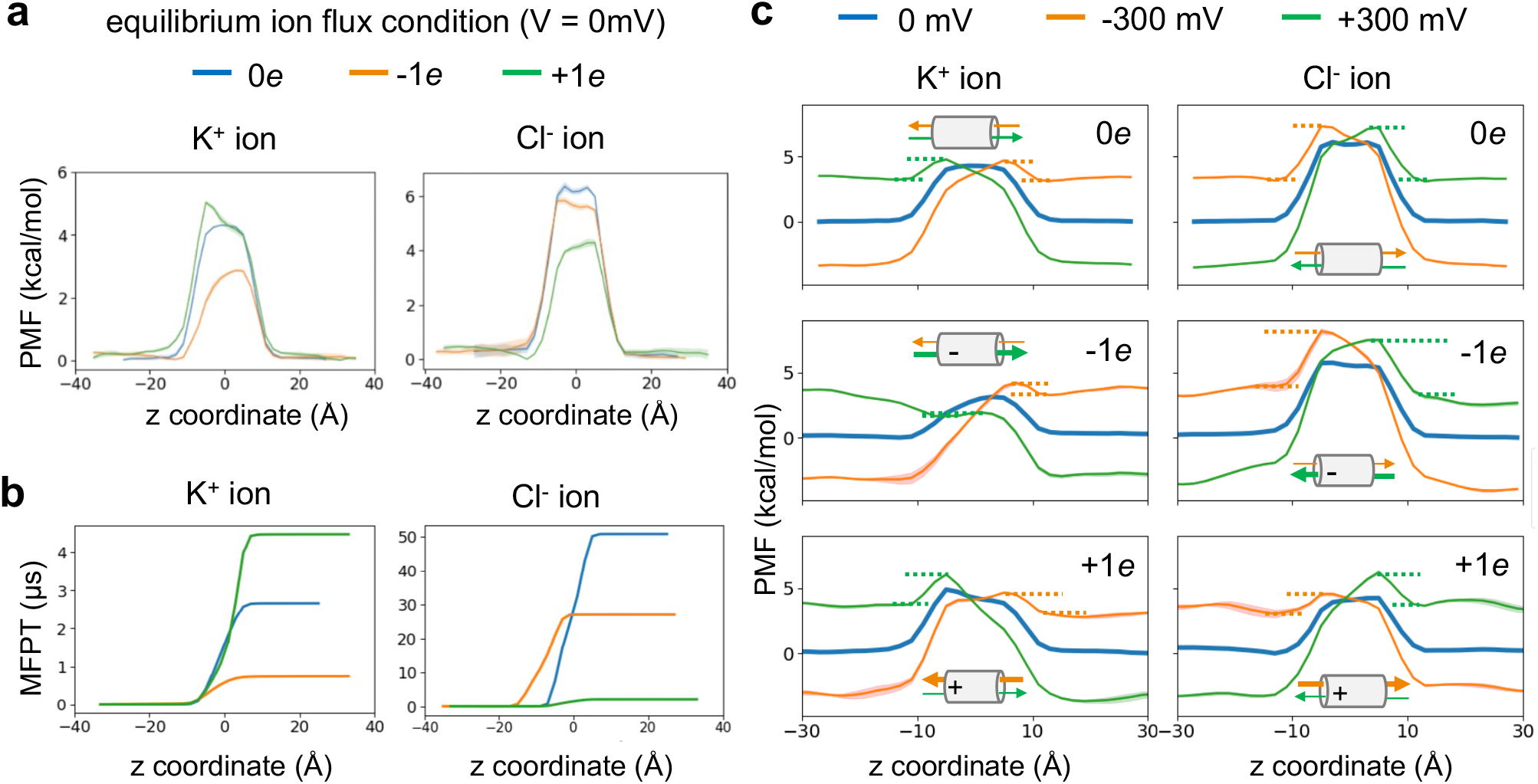
Free energy and kinetics of ion permeation in absence and presence of voltage. Equilibrium PMFs (**a**) and MFPTs (**b**) calculated for both K^+^ and Cl^-^ ions through each nanopore in absence of external driving force. (**c**) Derived PMFs in absence (blue) or presence of -300 mV (orange) or +300 mV (green) for each ion-nanopore system. Dotted lines represent permeation energy barriers. Shaded areas in PMF represent standard deviations obtained from dividing each Milestoning trajectory into three blocks. Rectification of net ion flux under voltage is qualitatively illustrated by thin vs. thick green and orange arrows in the nanopore cartoons. See also **Fig. S2-S4** for ± 600 mV data.

The presence of the intracellular -1*e* charge reduces the permeation energy barrier for both ions, but more so for potassium (4.1 to 2.8 kcal mol^-1^) than for chloride (6.0 to 5.5 kcal mol^-1^) ions (**Fig. 2a** orange), making the negative nanopore even more K^+^ selective compared to the neutral one. In contrast, the presence of the intracellular +1*e* charge slightly increases permeation energy barrier for K^+^ ions (4.1 to 5.0 kcal mol^-1^), but greatly reduces it for Cl^-^ ions (6.0 to 4.5 kcal mol^-1^) (**Fig. 2a** green), hence rendering the positive nanopore slightly more selective for Cl^-^ ions. Although these results are overall consistent with selectivity and relative conductance observed from voltage simulations, they do not tell us how voltage influences these processes and how nanopores rectify currents. To understand rectification, the effect of membrane potential on the intrinsic PMF must also be considered.

Since the carbon nanopore in our simulation systems is inherently rigid, only ions and water dipoles are under the influence of applied voltage. In this case, the PMF under voltage *W*_*tot*_(*z*) can be rigorously computed as the sum of the equilibrium PMF, *W*_*eq*_(*z*), and the additional potential introduced by the external field, *q*δ*ϕ*(*z*) (32-34). This additional potential has two components. The first one is the constant electric field throughout the entire simulated periodic cell, *E= V*(*z*)*/L*_*Z*_, where *V*(*z*) is the voltage linear to *L*_*Z*_, the length of the simulated system in the *z*-direction. The second is the reaction potential due to voltage-induced changes of the spatial distribution and orientation of water dipoles and ions (35). The reaction potential introduced by the external electrical field equates the difference in electrostatic potential in presence and absence of the external potential, *ϕ*_*v*_(*z*) − *ϕ*_0_(*z*). To calculate this difference, additional voltage simulations were carried out without ions inside the pore. The total PMF under each voltage, *W*_*tot*_(*z*), was obtained from **Eq. 1** (illustrated on **Fig. 3**) at +/-300 and +/-600 mV (**Fig. S2-S4**):

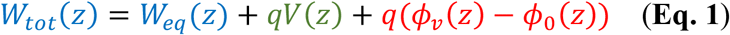

Because the equilibrium PMF of the neutral nanopore is symmetric for both ions, the leftward or rightward tilt of the PMF induced by positive or negative voltage is also symmetric, resulting in identical permeation energy barriers for inward and outward ionic permeation (1.5 and 4.2 kcal mol^-1^ for K^+^ and Cl^-^ ions, respectively) (**Fig. 2c** and **Table 1**). These symmetrical rearrangements of the PMF under voltage are consistent with the near-unity rectification score observed for potassium currents in the neutral nanopore (r = 0.9 ± 0.6) (**Table 1**). Due to the low chloride permeability of this nanopore, not a single Cl^-^ ion crossed it at +300 mV, whereas only one Cl^-^ ion crossed it at -300 mV during two independent 100 ns-long simulations. Hence, although we could not determine a chloride rectification score for the neutral nanopore, we expect it to also be near unity.

**Figure 3.**
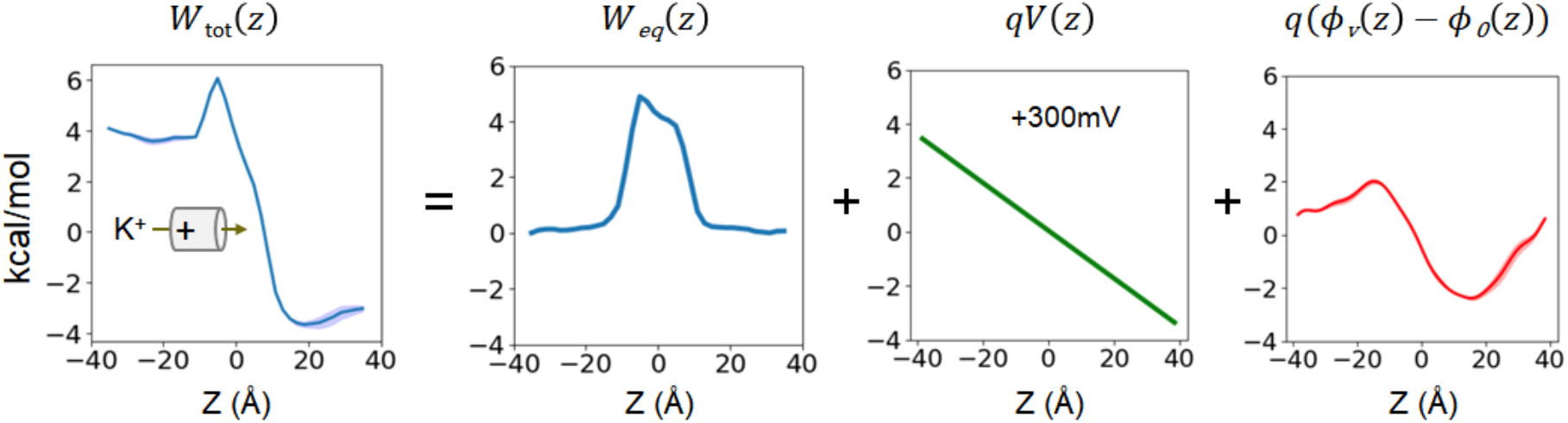
Example of PMF reconstruction for a K^+^ ion permeating through the positive nanopore at +300 mV. See **Fig. S2-S4** for all systems. Shaded area in red represents the standard deviation of the reaction field computed from two replicas of voltage simulations. The shaded area in total PMF is computed using error propagation.

In contrast to the neutral nanopore, both charged nanopores display an asymmetric equilibrium PMF. In these systems, the intracellular side of the PMF becomes shallower when the charge appended to this side attracts the permeating ion, and steeper when the appended charge repels it (**Fig. 2c**). This asymmetry causes a differential rearrangement of the energy barrier under voltage. Indeed, although the presence of a voltage accelerates ion permeation by decreasing the energy barrier in both directions, the degree to which this barrier decreases depends on the direction of the driving force: the permeation energy barrier under voltage is systematically higher when the driving force compel ions to enter the pore from the steeper side of the equilibrium PMF, and lower when ions enter through its shallower side (**Fig. 2c**). Hence, these PMF profiles under voltage fully recapitulate the rectification behavior observed in our voltage-driven flux simulations for both charged nanopores (**Fig. 1b**). In other words, PMF asymmetricity dictates how much the external driving force reduces permeation barrier depending on the direction to which it is applied, causing current rectification.

The additive nature of the potential of mean force under voltage (**Eq. 1**) tells us that the degree of rectification should also depend on the height of the asymmetric equilibrium PMF barrier relative to the magnitude of applied voltage. Indeed, when the gradient of voltage drop is much larger than the height of *W*_*eq*_(*z*), the shape of *W*_*tot*_(*z*) is dominated by *q*δ*ϕ*(*z*) to an extent that the asymmetrcity of *W*_*eq*_(*z*) becomes negligible. For example, when applying ±600 mV to the -1*e* nanopore, both inward and outward potassium permeation energy barriers vanish, rendering permeation nearly diffusion-limited for this ion (**Fig. S3**). Thus, one would expect that this type of current rectification becomes less prominent at higher voltage, as is sometimes observed across ion channel studies (36-38).

## Discussion

MD simulation, as an computational experiment, is a powerful tool to simulate current flow through ion channels. Although current rectification has been reported from MD simulations over two decades, a direct link between the observed rectified current and underlying asymmetric free energy profile were not investigated. In this study, we used all-atom molecular dynamics to illustrate and quantify how the intrinsic asymmetry of ion channels rectifies ionic currents, without the intervention of pore-blocking solutes or voltage-dependent structural rearrangements. By deconstructing the minimal physical requirement for rectification, we establish a quantitative relationship between free energy profile and ion-specific rectification rate.

The microscopic permeation kinetics of ion crossing a channel under voltage is known to be related to the rate coefficient, *k*, calculated according to Eyring’s rate theory (39):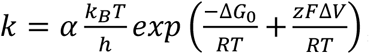, where 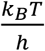 is the vibration frequency under thermal energy, with *k*_*B*_ the Boltzmann’s constant (1.38 × 10^−23^ J K^-1^), *T* the temperature (here 300.15 K) and *h* the Planck’s constant (6.626 × 10^−34^ J s); Δ*G*_0_ (kcal mol^-1^) the energy barrier without voltage from *W*_*eq*_(*z*); *R* the gas constant (8.314 J mol^-1^); *z* the charge; *F* the Faraday constant, Δ*V* the portion of voltage gradient affecting barrier crossing, and α, a dimensionless constant (sometimes called transmission coefficient). However, this equation is inherently unable to predict a rectification behavior arising from a single energy barrier, because the height of this energy barrier, Δ*G*_0_, is determined under equilibrium, which implies that Δ*G*_0_ has the same value in both permeation directions. This is why the Eyring model requires at least a binding site separated by two energy barriers of different height to predict rectification (22).

Here, because we have obtained the energy barrier under voltage, Δ*G*_*v*_, directly from *W*_*tot*_(*z*), the apparent rate coefficient of ion crossing the barrier under voltage, 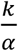, can be directly calculated from our data as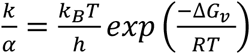. The calculated 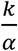 values reproduce outward rectification observed in the negative nanopore (r = 2.30 and 1.40 for K^+^ and Cl^-^ ions, respectively), and inward rectification in the positive nanopore (r = 0.22 and 0.11 for K^+^ and Cl^-^ ions, respectively) (**Table S1**). Across all ion-nanopore systems, the relative conductance values estimated from *W*_*tot*_(*z*) correlate well with the relative conductance values obtained from MD simulations (proportionality coefficient = 1.03 ± 0.08, R^2^ > 0.97) (**Fig. S5**). Such a quantitative agreement between PMF barrier and voltage-driven flux further confirms that the asymmetric PMF derived from MD simulations can be used to understand and predict the degree of the rectification for a specific ion.

The asymmetric free energy profiles generated in this study bare certain similarities to Feynman’s ratchet model for the steady-state directed motion (40), where the ratchet potential is shaped like an asymmetric sawtooth. However, the ratchet potential includes a binding site, which we show here is not necessary for rectification. While we used electrical charges to generate PMF asymmetry, other factors such as the presence of ion binding sites or asymmetric pore geometry could also contribute to PMF asymmetry. Indeed, most ion channel pores are far from perfect cylinders, and many ion channels harbor ion binding sites within their pores. Yet, experimental (23, 24) and computational (15) studies have shown that geometric asymmetry is not a sufficient condition for rectification unless an asymmetric electrostatic potential is also present, which is the focus of this study. In addition, the presence of ion binding sites may contribute to rectification but only if they induce PMF asymmetry, such as when the two adjacent barriers flanking them have different heights (22). In conclusion, our voltage-dependent PMFs demonstrate that rectification caused by energy barrier asymmetry depends on the nature of the permeating ion, can be tuned by electrostatic pore polarity, does not require ion binding sites, conformational flexibility, or specific pore geometry, and, as such, may be widespread among ion channels.

Rectification caused by PMF asymmetry can be generalized to other types of steady-state flux. In this study, we applied an external electrical field to bias ion permeation toward a specific direction, showing that the interaction of this external driving force with an asymmetrically-shaped PMF results in different inward and outward current magnitude. In principle, rectification could also arise from the interaction of an asymmetric PMF with another source of external driving forces, such as concentration gradients. Therefore, this work provides a unified approach to dissect intrinsic current rectification in depth and can be broadly applied to the fields of ion channel and material sciences.

## Acknowledgments

This work was supported by NIH grant GM130834 to J.J.L. and Y.L.L.

## Authors Contributions

Project conception: Y.L., J.D.S., J.J.L. and Y.L.L., Data acquisition and analyses: Y.L. and Y.L.L. Manuscript writing: J.J.L. and Y.L.L. with inputs from all authors.

## Supplementary information

**Table S1.**
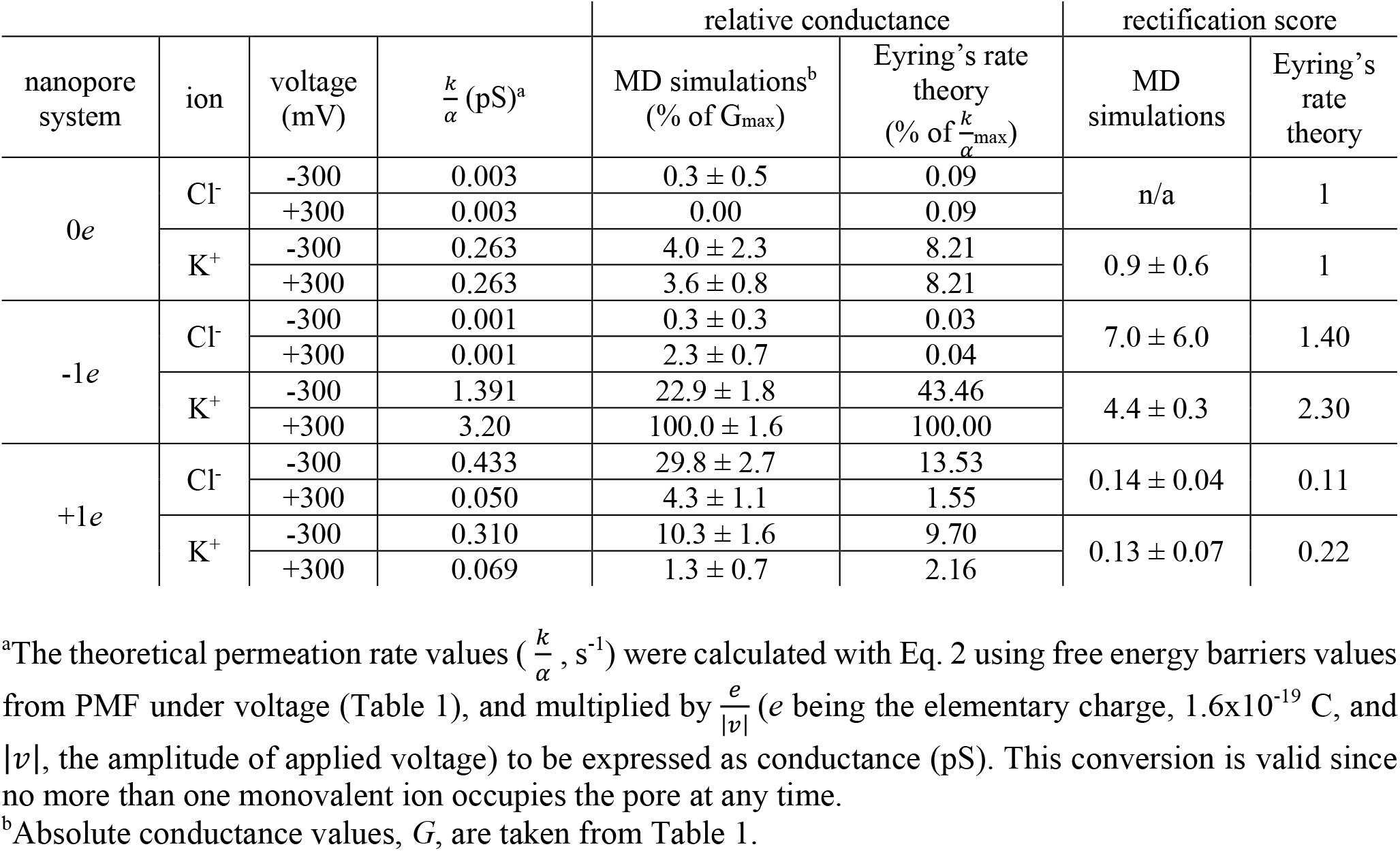
Conductance and rectification data determined from MD simulations and from Eyring’s rate theory.

**Figure S1.**
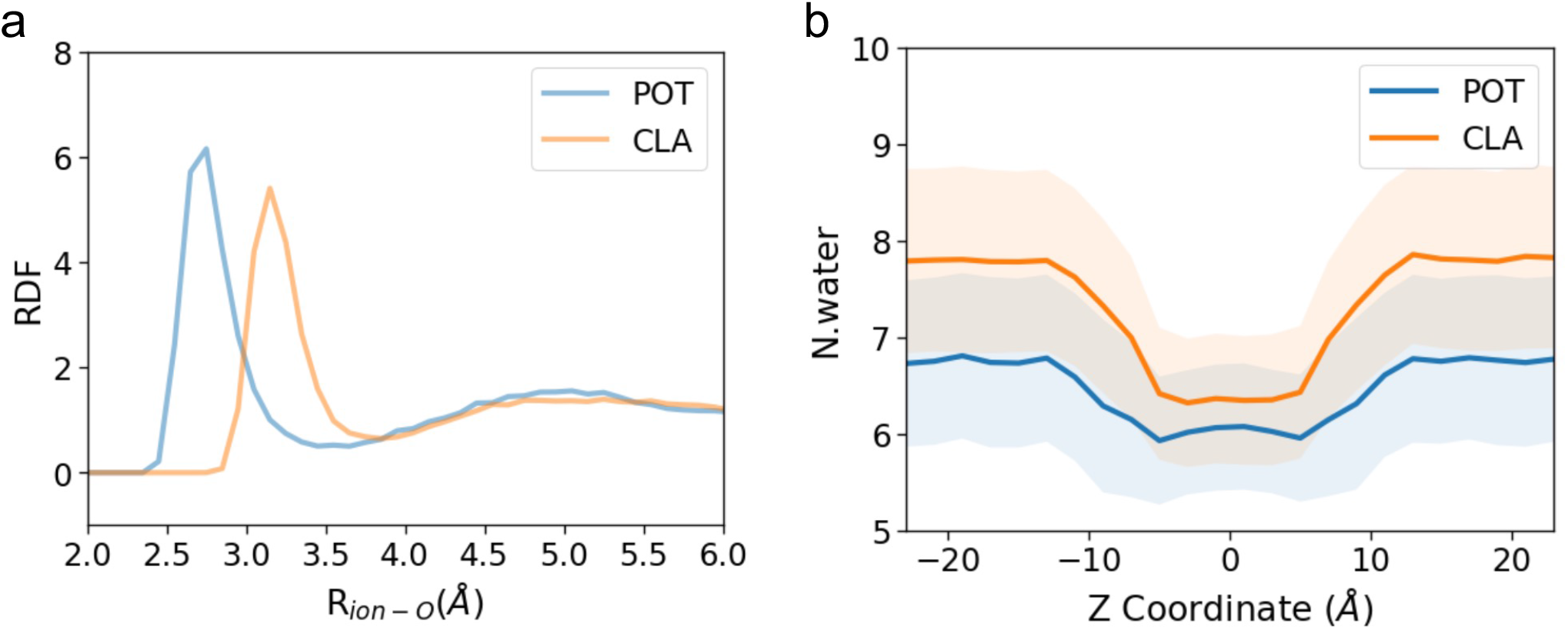
**a**. Radial distribution function (RDF) of ion-water oxygen in bulk, averaged over 8000 frames (80 ns). The hydration radii of Cl^-^ and K^+^ (3.85 Å and 3.45 Å) correspond to the location of the first minimum. **b**. The change in the number of water oxygen within the hydration radii of Cl^-^ and K^+^ along the z-axis of the neutral nanopore system, computed from each Voronoi cell of the milestoning simulations. The shaded area represents the standard deviation from the 5000 frames (50ns).

**Figure S2.**
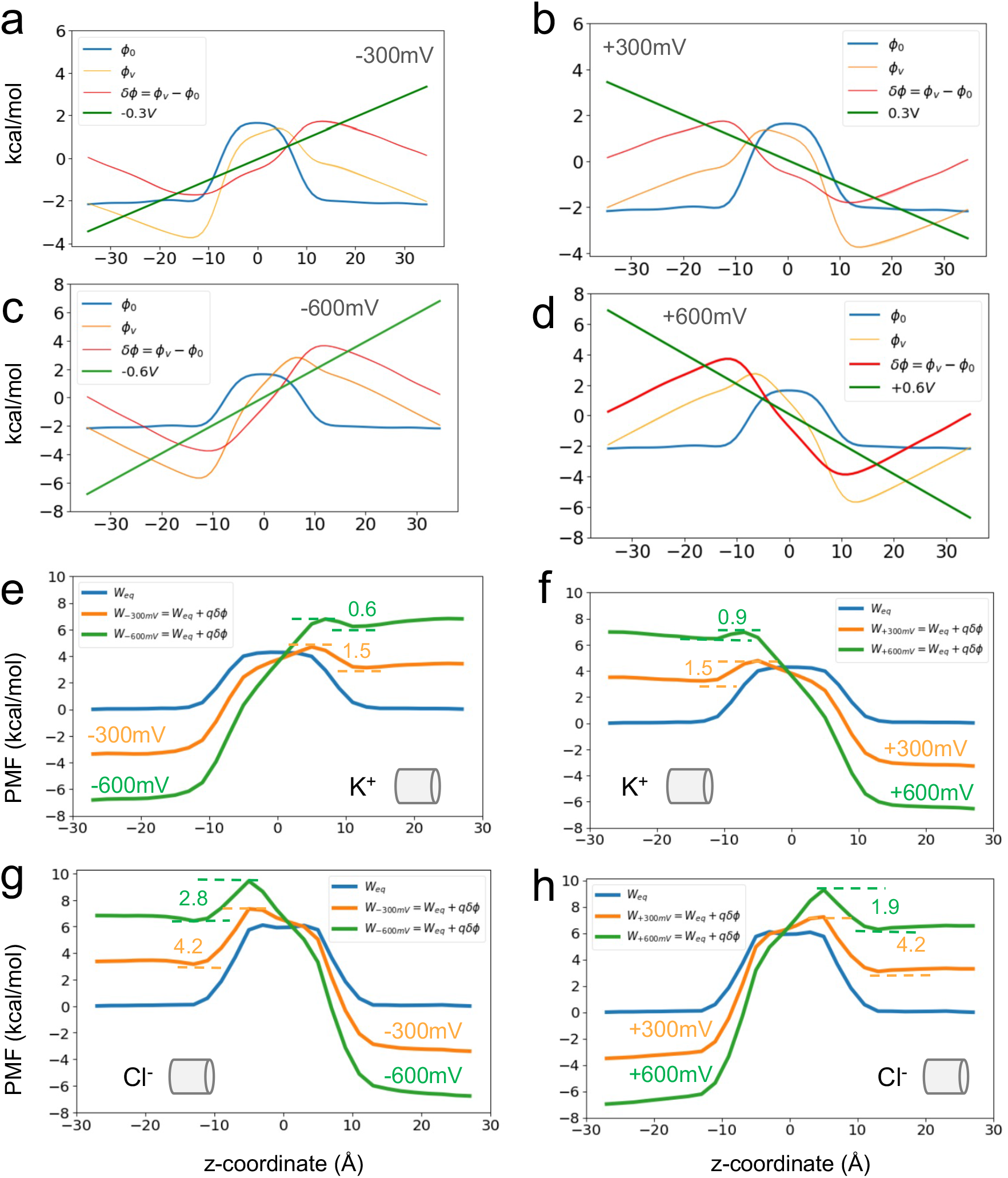
Reconstruction of the permeation PMF under voltage for the neutral nanopore. **(a-d)** deconstruction of additional potential into the constant electric field (green) and the reaction potential in absence (blue) and presence (yellow) of external voltage. (**e-h**) Reconstructed PMF for each ion and voltage. Numbers in plots indicate energy barrier values in kcal/mol.

**Figure S3.**
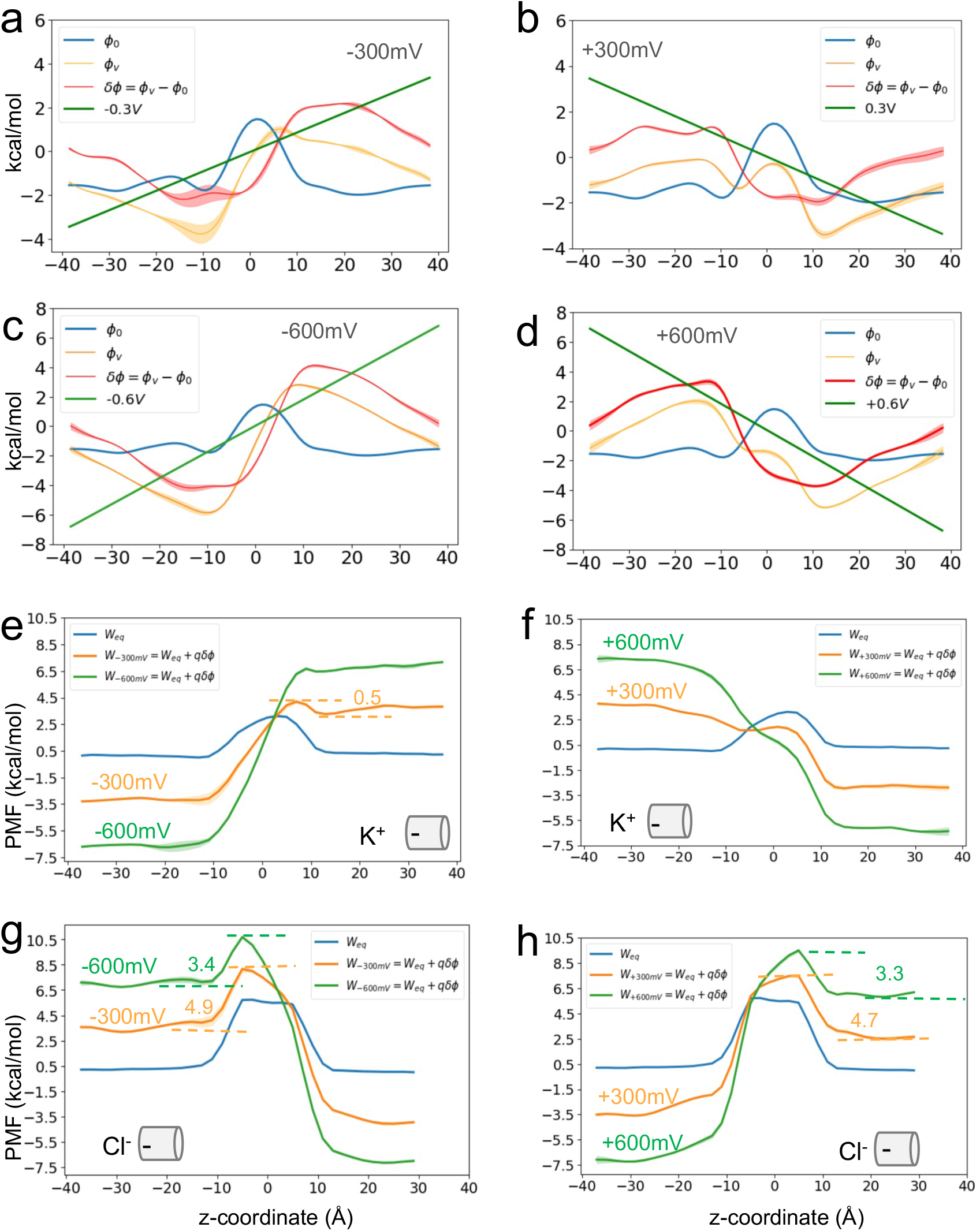
Reconstruction of the permeation PMF under voltage for the negative nanopore. **(a-d)** deconstruction of additional potential into the constant electric field (green) and the reaction potential in absence (blue) and presence (yellow) of external voltage. (**e-h**) Reconstructed PMF for each ion and voltage. Numbers in plots indicate energy barrier values in kcal/mol.

**Figure S4.**
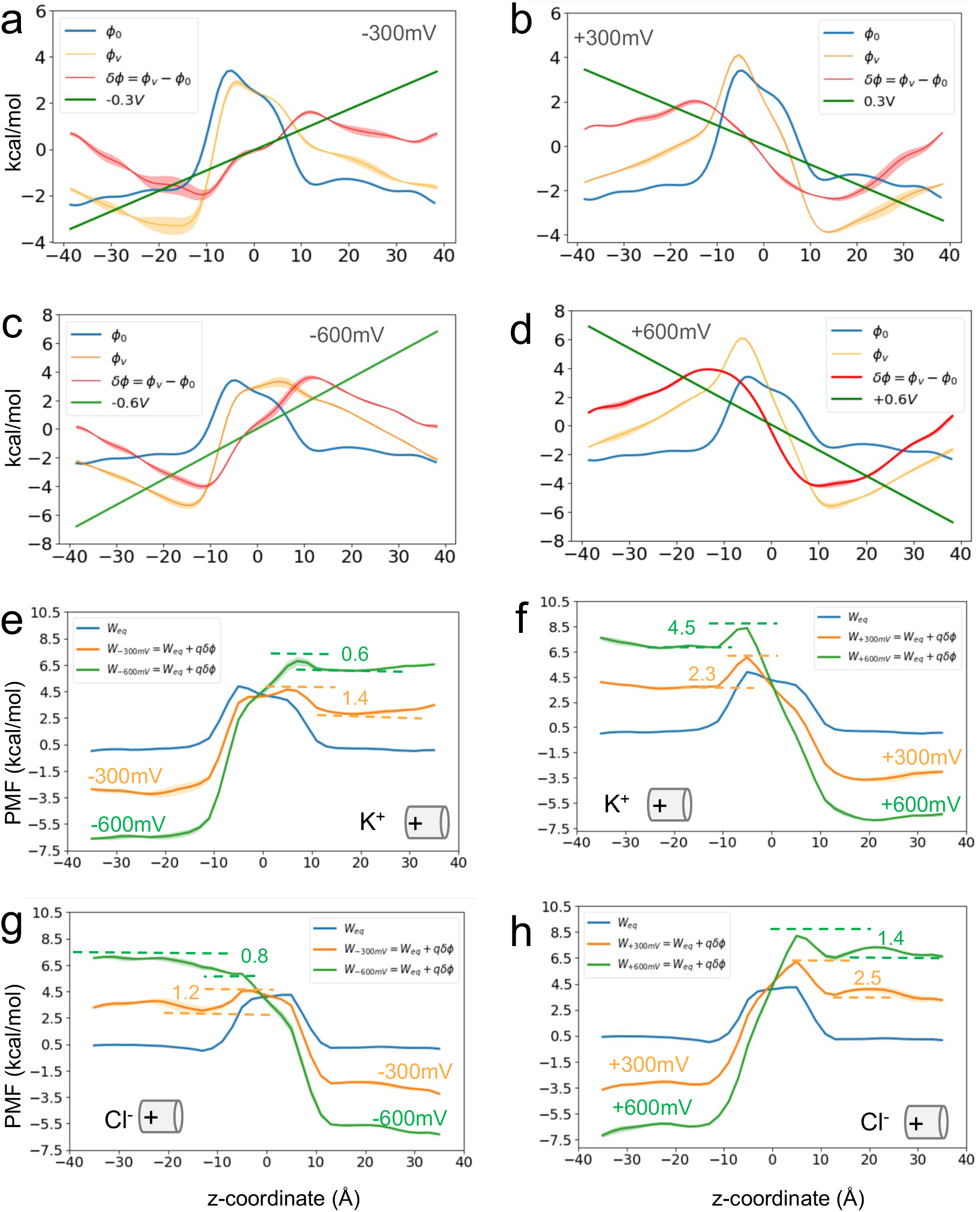
Reconstruction of the permeation PMF under voltage for the positive nanopore. **(a-d)** deconstruction of additional potential into the constant electric field (green) and the reaction potential in absence (blue) and presence (yellow) of external voltage. (**e-h**) Reconstructed PMF for each ion and voltage. Numbers in plots indicate energy barrier values in kcal/mol.

**Figure S5.**
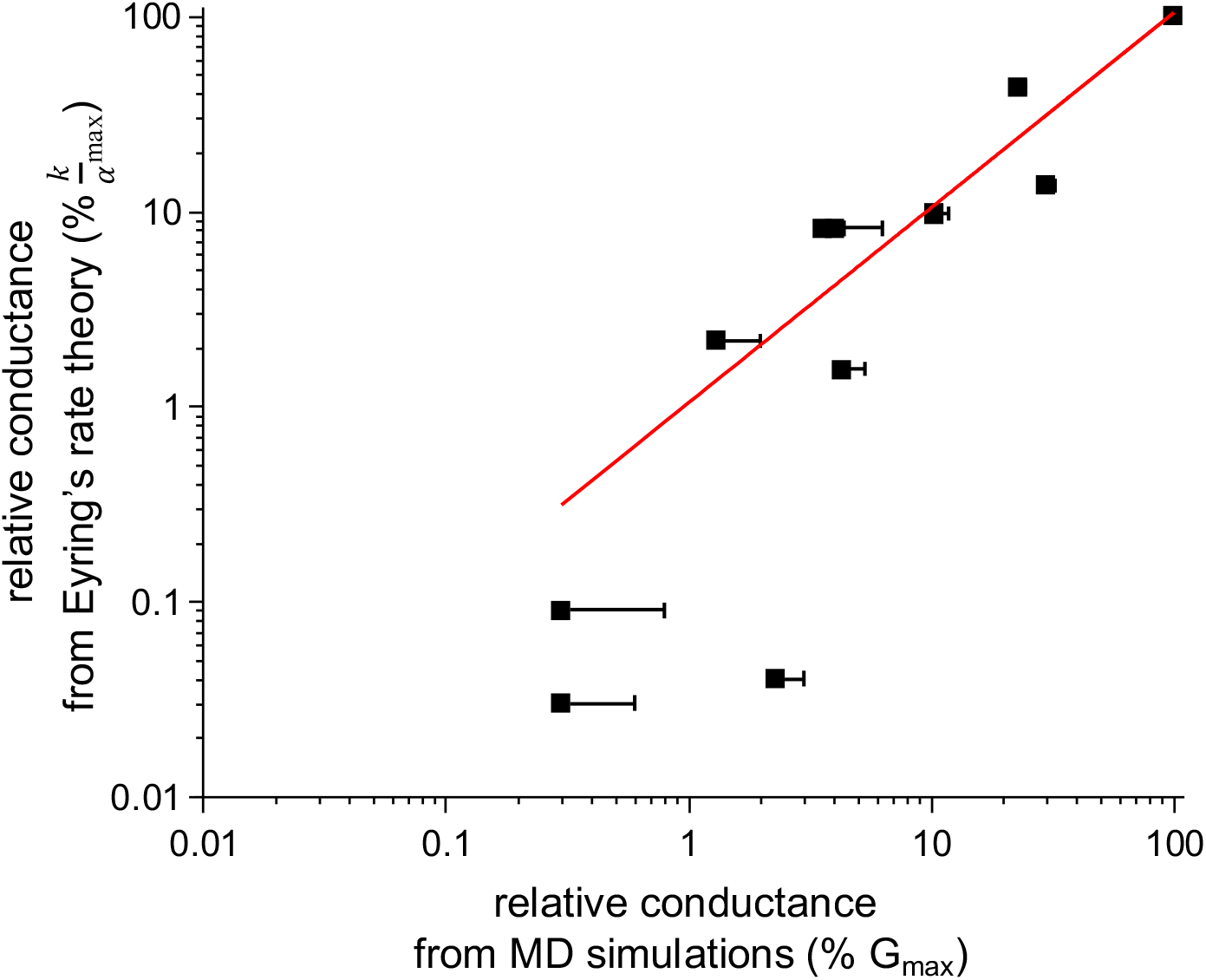
Comparison of conductance determined from MD simulations and from Eyring’s rate theory. Relative conductance values determined from Eyring’s rate theory were plotted against those measured from MD simulations (see Table S1). Fitting the data using the linear equation *y* = *bx* returned a proportionality constant near unity (b = 1.03 ± 0.08, R^2^ = 0.97952, red line). The data are shown in a log scale for clarity. Error bars in the x-axis are shown as positive standard deviations.

## References

1. Daiguji, H., Y. Oka, and K. Shirono. 2005. Nanofluidic diode and bipolar transistor. Nano Lett 5(11):2274–2280.

2. Karnik, R., C. Duan, K. Castelino, H. Daiguji, and A. Majumdar. 2007. Rectification of ionic current in a nanofluidic diode. Nano Lett 7(3):547–551.

3. Vlassiouk, I., and Z. S. Siwy. 2007. Nanofluidic diode. Nano Lett 7(3):552–556.

4. Zhang, Y., and G. C. Schatz. 2017. Advantages of Conical Pores for Ion Pumps. J Phys Chem C 121(1):161–168.

5. Lu, Z. 2004. Mechanism of rectification in inward-rectifier K+ channels. Annu Rev Physiol 66:103–129.

6. Bezanilla, F. 2000. The voltage sensor in voltage-dependent ion channels. Physiol Rev 80(2):555–592.

7. Nilius, B., J. Prenen, G. Droogmans, T. Voets, R. Vennekens, M. Freichel, U. Wissenbach, and V. Flockerzi. 2003. Voltage dependence of the Ca2+-activated cation channel TRPM4. J Biol Chem 278(33):30813–30820.

8. Lewis, A., Z. A. McCrossan, R. W. Manville, M. O. Popa, L. G. Cuello, and S. A. N. Goldstein. 2020. TOK channels use the two gates in classical K(+) channels to achieve outward rectification. FASEB J 34(7):8902–8919.

9. Busath, D., and G. Szabo. 1981. Gramicidin forms multi-state rectifying channels. Nature 294(5839):371–373.

10. Overholt, J. L., A. Saulino, M. L. Drumm, and R. D. Harvey. 1995. Rectification of whole cell cystic fibrosis transmembrane conductance regulator chloride current. Am J Physiol 268(3 Pt 1):C636–646.

11. Martins, J. R., D. Faria, P. Kongsuphol, B. Reisch, R. Schreiber, and K. Kunzelmann. 2011. Anoctamin 6 is an essential component of the outwardly rectifying chloride channel. P Natl Acad Sci USA 108(44):18168–18172.

12. Luo, Y., B. Egwolf, D. E. Walters, and B. Roux. 2010. Ion selectivity of alpha-hemolysin with a beta-cyclodextrin adapter. I. Single ion potential of mean force and diffusion coefficient. J Phys Chem B 114(2):952–958.

13. Dessaux, D., J. Mathe, R. Ramirez, and N. Basdevant. 2022. Current Rectification and Ionic Selectivity of alpha-Hemolysin: Coarse-Grained Molecular Dynamics Simulations. J Phys Chem B.

14. Bhattacharya, S., L. Muzard, L. Payet, J. Mathe, U. Bockelmann, A. Aksimentiev, and V. Viasnoff. 2011. Rectification of the current in alpha-hemolysin pore depends on the cation type: the alkali series probed by MD simulations and experiments. J Phys Chem C Nanomater Interfaces 115(10):4255–4264.

15. Liu, Y., and F. Zhu. 2013. Collective diffusion model for ion conduction through microscopic channels. Biophys J 104(2):368–376.

16. Constantin, D., and Z. S. Siwy. 2007. Poisson-Nernst-Planck model of ion current rectification through a nanofluidic diode. Phys Rev E Stat Nonlin Soj Matter Phys 76(4 Pt 1):041202.

17. Tang, L., Y. Hao, L. Peng, R. Liu, Y. Zhou, and J. Li. 2024. Ion current rectification properties of non-Newtonian fluids in conical nanochannels. Phys Chem Chem Phys 26(4):2895–2906.

18. Trivedi, M., and N. Nirmalkar. 2022. Ion transport and current rectification in a charged conical nanopore filled with viscoelastic fluids. Sci Rep 12(1):2547.

19. van Oeffelen, L., W. Van Roy, H. Idrissi, D. Charlier, L. Lagae, and G. Borghs. 2015. Ion current rectification, limiting and overlimiting conductances in nanopores. PLoS One 10(5):e0124171.

20. Cruz-Chu, E. R. a. A., Aleksei and Schulten, Klaus. 2009. Ionic current rectification through silica nanopores. The Journal of Physical Chemistry C 113:1850--1862.

21. Lan, W. J., M. A. Edwards, L. Luo, R. T. Perera, X. Wu, C. R. Martin, and H. S. White. 2016. Voltage-Rectified Current and Fluid Flow in Conical Nanopores. Acc Chem Res 49(11):2605–2613.

22. Woodbury, J. W. 1969. Eyring Rate Theory Model of the Current-Voltage Relationships of Ion Channels in Excitable Membranes. Advance in Chemical Physics. I. Prigogine and S. A. Rice, editors. John Wiley & Sons Ltd.

23. Siwy, Z., E. Heins, C. C. Harrell, P. Kohli, and C. R. Martin. 2004. Conical-nanotube ion-current rectifiers: the role of surface charge. J Am Chem Soc 126(35):10850–10851.

24. Siwy, Z. S. 2006. Ion-current rectification in nanopores and nanotubes with broken symmetry. Adv Funct Mater 16(6):735–746.

25. Kosinska, I. D. 2006. How the asymmetry of internal potential influences the shape of I-V characteristic of nanochannels. J Chem Phys 124(24):244707.

26. Lin, Y. C., and Y. L. Luo. 2022. Unifying Single-Channel Permeability From Rare-Event Sampling and Steady-State Flux. Front Mol Biosci 9:860933.

27. MacKerell Jr, A. D., D. Bashford, M. Bellott, R. L. Dunbrack Jr, J. D. Evanseck, M. J. Field, S. Fischer, J. Gao, H. Guo, S. Ha, D. Joseph-McCarthy, L. Kuchnir, K. Kuczera, T. K. Lau, C. Mattos, S. Michnick, T. Ngo, D. T. Nguyen, R. Stote, J. Straub, M. Watanabe, J. Wiorkiewicz-Kuczera, D. Yin, and M. Karplus. 1998. All-atom empirical potential for molecular modeling and dynamics studies of proteins. J Phys Chem B 102(18):3586–3616.

28. Mackerell Jr, A. D., M. Feig, and C. L. Brooks III. 2004. Extending the treatment of backbone energetics in protein force fields: Limitations of gas-phase quantum mechanics in reproducing protein conformational distributions in molecular dynamics simulations. J Comput Chem 25(11):1400–1415.

29. Darden, T., D. York, and L. Pedersen. 1993. Particle mesh Ewald: An N·log (N) method for Ewald sums in large systems. J Chem Phys 98(12):10089–10092.

30. Aksimentiev, A., and K. Schulten. 2005. Imaging alpha-hemolysin with molecular dynamics: Ionic conductance, osmotic permeability, and the electrostatic potential map. Biophysical Journal 88(6):3745–3761.

31. Jiang, W., Y. C. Lin, W. Botello-Smith, J. E. Contreras, A. L. Harris, L. Maragliano, and Y. L. Luo. 2021. Free energy and kinetics of cAMP permeation through connexin26 via applied voltage and milestoning. Biophys J 120(15):2969–2983.

32. Berneche, S., and B. Roux. 2003. A microscopic view of ion conduction through the K+ channel. Proc Natl Acad Sci 100(15):8644–8648.

33. Roux, B. 1999. Statistical mechanical equilibrium theory of selective ion channels. Biophys J 77(1):139–153.

34. Roux, B. 2008. The membrane potential and its representation by a constant electric field in computer simulations. Biophys J 95(9):4205–4216.

35. Gumbart, J., F. Khalili-Araghi, M. Sotomayor, and B. Roux. 2012. Constant electric field simulations of the membrane potential illustrated with simple systems. Biochim Biophys Acta 1818(2):294–302.

36. Numata, T., T. Shimizu, and Y. Okada. 2007. TRPM7 is a stretch- and swelling-activated cation channel involved in volume regulation in human epithelial cells. Am J Physiol-Cell Ph 292(1):C460–C467.

37. Chung, M. K., A. D. Guler, and M. J. Caterina. 2005. Biphasic currents evoked by chemical or thermal activation of the heat-gated ion channel, TRPV3. J Biol Chem 280(16):15928–15941.

38. Miller, D. R., H. Khoshbouei, S. Garai, L. N. Cantwell, C. Stokes, G. Thakur, and R. L. Papke. 2020. Allosterically Potentiated alpha7 Nicotinic Acetylcholine Receptors: Reduced Calcium 1. Permeability and Current-Independent Control of Intracellular Calcium. Mol Pharmacol 98(6):695–709.

39. Hille, B. 2001. Ion channels of excitable membranes. Sinauer, Sunderland, Mass.

40. Feynman, R. P., R. B. Leighton, and M. Sands. 1963. Ratchet and Pawl. The Feynman Lectures on Physics, pp. 46–51.

